# Computational design of Matrix Metalloprotenaise-9 (MMP-9) resistant to auto-cleavage

**DOI:** 10.1101/2023.04.11.536383

**Authors:** Alessandro Bonadio, Solomon Oguche, Tali Lavy, Oded Kleifeld, Julia Shifman

**Author notes:** These authors contributed equally (Alessandro Bonadio, Solomon Oguche).

## Abstract

Matrix metalloproteinase-9 (MMP-9) is an endopeptidase that remodels the extracellular matrix and has been implicated as a major driver in cancer metastasis. Hence, there is a high demand for MMP-9 inhibitors for therapeutic purposes. For such drug design efforts, large amounts of MMP-9 are required. Yet, the catalytic domain of MMP-9 (MMP-9_Cat_) is an intrinsically unstable enzyme that tends to auto-cleave within minutes, making it difficult to use in drug design experiments and other biophysical studies. We set our goal to design MMP-9_Cat_ variant that is active but stable to autocleavage. For this purpose, we first identified potential autocleavage sites on MMP-9_Cat_ using mass spectroscopy and then eliminated the autocleavage site by predicting mutations that minimize autocleavage potential without reducing enzyme stability. Four computationally designed MMP-9_Cat_ variants were experimentally constructed and evaluated for auto-cleavage and enzyme activity. Our best variant, Des2, with 2 mutations, was as active as the wild-type enzyme but did not exhibit auto-cleavage after seven days of incubation at 37°C. This MMP-9_Cat_ variant, with an identical to MMP- 9_Cat_ WT active site, is an ideal candidate for drug design experiments targeting MMP-9 and enzyme crystallization experiments. The developed strategy for MMP-9_CAT_ stabilization could be applied to redesign of other proteases to improve their stability for various biotechnological applications.

## Introduction

Matrix metalloproteinases (MMPs) are a family of proteases, comprising more than twenty different enzymes in humans. They are composed of a catalytic domain with a catalytic zinc ion, a pro-domain which is cleaved off upon enzyme activation by MMPs and other proteases [1,2], an optional transmembrane domain, and additional domains utilized in substrate and other protein binding [3]. MMPs are involved in multiple biological processes, including remodeling of the extracellular matrix (ECM), shedding of cell surface receptors and membrane-bound signaling molecules, angiogenesis, intravasation/extravasation from blood vessels, and immune cells maturation [4]. It is not surprising that MMPs play an important role in cancer metastasis, with multiple MMPs overexpressed in solid tumors [4–9]. Due to the important role of MMPs in cancer, there has been considerable effort to develop MMP inhibitors, from zinc chelators [10–12] to antibodies [13–17]. In particular, several drug design efforts have been directed against MMP-9. A monoclonal antibody Andecaliximab directed against MMP-9 is being evaluated in multiple phase 3 clinical trials involving solid tumors [15,16]. We and others have engineered potent and selective inhibitors of MMP-9 starting from the endogenous MMP inhibitor N-TIMP2 [18–22].

Drug design efforts targeting MMP-9 require large amounts of the enzyme being produced. However, working with MMP-9 has been challenging due to enzyme instability. Indeed, all routine procedures such as protein labeling, dialysis, crystallization, and other experiments requiring long incubations and high protein concentrations are extremely challenging with this enzyme due to the fast loss of the active protein. The major reason for this loss is the capability of MMP-9 to auto-degrade; self-cleavage becomes more prominent at higher protein concentrations [23,24]. To obviate the problem, weak MMP inhibitors such as acetohydroxamic acid have been used in MMP-9 purification procedures. However, the inhibitor has to be removed prior to further experiments and upon removal, the self-cleavage problem returns [25]. To alleviate self-cleavage and improve MMP-9 stability, MMP-9 mutants that remove the catalytic Glu and abolish catalytic activity, E402Q and E402A, have been utilized in some studies [25,26]. Such enzyme variants are more stable but present a modified active site, with the negative charge of E402 removed. Thus, these active-site MMP-9 mutants could not be used in MMP-9 drug design or screening experiments.

In the current study, we present a novel approach for engineering the active catalytic domain of MMP-9 variants that are resistant to self-cleavage by identifying and removing the self-cleavage site from the enzyme sequence. Using computational design, we introduce a minimum number of mutations that preserve protein structure and activity but decrease the probability of sequence cleavage by MMP-9. Our best design with two mutations 10 Å away from the active site does not auto-cleave after seven days of incubation at 37 C while retaining WT-like MMP-9 activity.

## Results

### Design of the non-degrading MMP-9 variants

For our studies, we utilized the MMP-9 construct that contained only the MMP-9 catalytic domain construct referred to as MMP-9_Cat_ (residues 107–215 fused to 391– 443) with the hemopexin-like and fibronectin type II domains truncated [26]. We have been routinely expressing MMP-9_Cat_ in *E. Coli* [20], refolding the enzyme from inclusion bodies, and performing subsequent purification with size exclusion chromatography (SEC). While this protocol always yielded active protein, more than 90% of the protein was lost during refolding and purification. The loss of active MMP-9_Cat_ WT was at least partially due to the fast self-cleavage of the enzyme, as could be observed on the SDS-PAGE gel analysis of the pure MMP-9_Cat_ WT sample (Fig. 1A).

**Figure 1.**
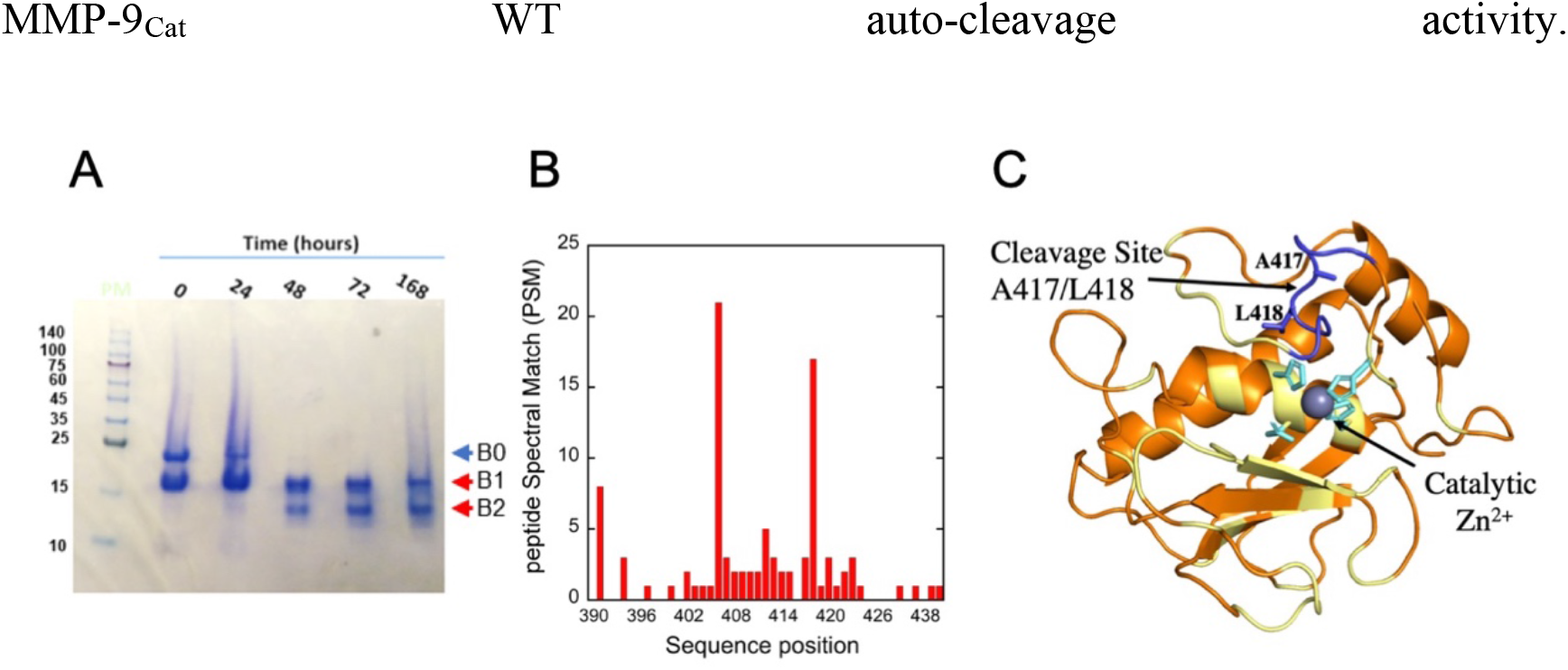
MMP-9 auto-degradation site is a partially exposed loop. (**A**) SDS-PAGE gel analysis of MMP-9_Cat_ WT at 25ºC showing the full length (blue arrow) and degradation products (red arrows). (**B**) The relative auto-cleavage preference of MMP-9_Cat_ WT. Cleavage preference was calculated from the LS/MS count of each peptide product contributing to a given cleavage site at a specific sequence position. The X-axis represents the amino acid residue number of the full MMP-9 sequence. The C-terminal cleavage positions with the highest Peptide Spectral Match (PSM) were after positions 405 and 417. (**C**) Structure of MMP-9_Cat_ WT (orange) in cartoon representation, showing the catalytic Zn^2+^ and the predicted auto-cleavage site in the partially exposed loop (colored in blue). Amino acids at the interface with the MMP protein inhibitor TIMP-2 are colored in yellow. Zn^2+^ ligating residues are colored in cyan.

To identify the MMP-9_Cat_ WT auto-cleavage site(s), we extracted the early auto-cleavage product (Band B1 on the gel), digested it with trypsin, and performed the Liquid Chromatography-Mass Spectroscopy (LC-MS) analysis (Supplementary data). Our data analysis aimed at the identification of semi-tryptic peptides whose non-tryptic-termini were generated due to MMP-9_Cat_ self-cleavage. This analysis showed four possible cleavage sites: the N-terminal sites with P1 positions 185 and 190 and the C-terminal sites with P1 positions 405 and 417 on MMP-9_Cat_ WT (Fig. 1B). Only the C-terminal cleavage sites, however, were consistent with the cleavage product of 16 kDa observed by SDS-PAGE (B1 on Fig. 1A). Among the two identified C-terminal cleavage sites, the cleavage site between positions A417 and L418 on MMP-9_Cat_ WT had been identified in a previous study that analyzed the MMP-9 self-cleavage profile [27]. Inspection of the A417/L418 self-cleavage site revealed that it is 10 Å away from the active site and away from the MMP-9 binding interface with the broad MMP protein inhibitor TIMP-2 and small molecule MMP inhibitors (Fig. 1C). We hence decided that the A417/L418 is the most promising site for redesign with the goal of eliminating MMP-9_Cat_ WT auto-cleavage activity.

We set our goal to design a stable to auto-cleavage MMP-9_Cat_ variant by introducing mutations in the auto-cleavage site that reduce the MMP-9_Cat_ auto-cleavage score but at the same time do not destabilize the enzyme. We wanted to introduce a minimal number of mutations in order to keep the MMP-9_Cat_ sequence and structure as similar to WT as possible, preserving the active site and the binding interface with a broad MMP family inhibitor TIMP-2. We hence considered four amino acids before and after the identified cleavage site, as commonly done in the analysis of protease substrate specificity [28]. We then ran a single-position mutational scan of these eight MMP-9_Cat_ positions and calculated the auto-cleavage score using the Procleave web server [29]. At the same time, protein stability of all single mutants was predicted using Rosetta FastDesign [30], thus generating both self-cleavage probability matrix and stability matrix for all possible positions at these eight MMP-9_Cat_ residues (Fig. 2A-B).

**Figure 2.**
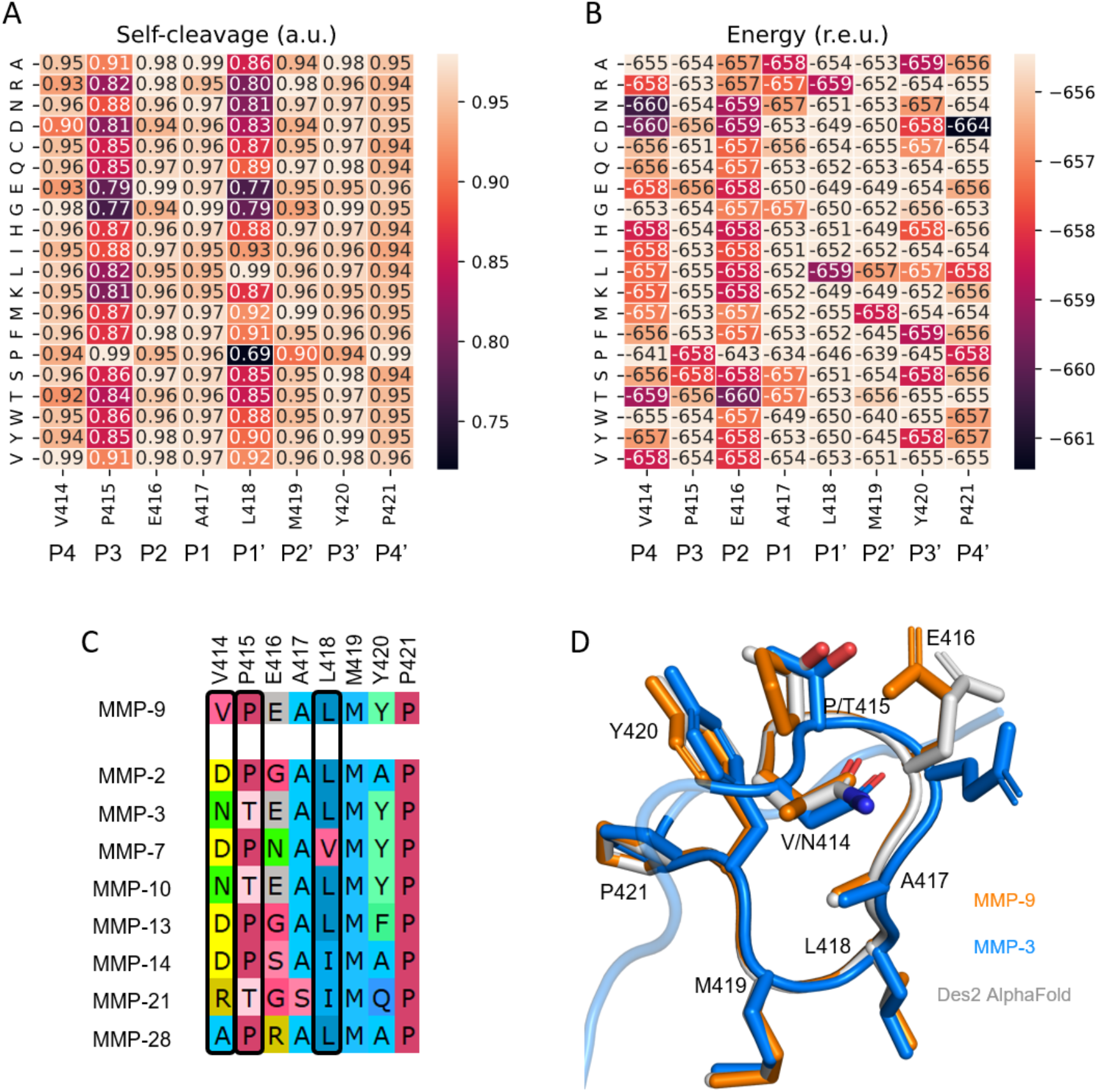
Design of non-degrading MMP-9_Cat_ variants. (**A**) Saturation mutagenesis scan of the cleavage loop displaying the auto-cleavage score according to the Procleave webserver [29]. (**B**) Saturation mutagenesis scan of the cleavage loop displaying the Rosetta energy. (**C**) Eight most similar to MMP-9 MMPs were aligned in the region of the predicted auto-cleavage site. D414, N414, T415, and I418 are frequently present in MMP-9 homologs and confer lower predicted auto-cleavage scores compared to that of the MMP-9_Cat_ WT sequence and exhibit favorable energy according to Rosetta. (**D**) A model of the Des2 loop according to AlphaFold (light gray) superimposed onto the structures of MMP-3 (blue, PDB: 4DP3) and MMP-9 (orange, PDB: 5TH9). The mutated N414 and T415 in Des2 have the same rotamers as in the homologous MMP-3 structure as well as the same neighboring amino acids and most of the same rotamers among neighboring positions, increasing the confidence in Des2 foldability.

We then compared the computational scans for auto-degradation and stability and the multiple sequence alignments of MMPs for those eight positions and selected mutations that improved self-cleavage score without compromising stability (Fig. 2C). Comparing the design models to the structure of the homologs, we further selected designs that recapitulated homologous structures, both at the mutated positions and its structural neighbors, producing Des1 and Des2 (Fig. 2D). A few additional mutations were introduced with RosettaFast design at other MMP-9_Cat_ positions to stabilize the designs further, producing Des3 and Des4 sequences. Thus, four MMP-9_Cat_ designs targeting different positions were selected for experimental characterization (Table 1).

**Table 1.**
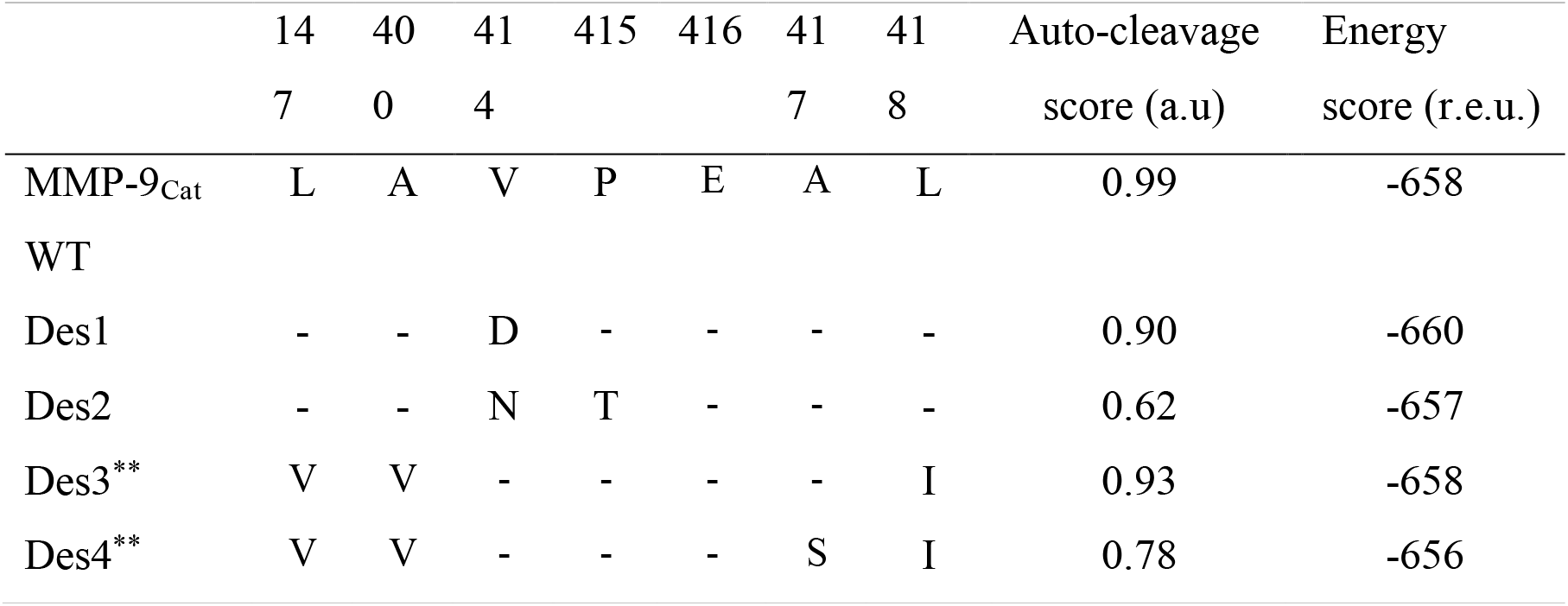
The four design sequences with the predicted auto-cleavage and energy scores. ^**^Des3 and Des4 have the additional buried mutations L147V and A400V, which greatly stabilize the core of the protein and are present in other homologs.

### Experimental characterization of the designed MMP-9_Cat_ variants

The MMP-9_Cat_ WT and four variants (Des1, Des2, Des3, and Des4) were expressed in *E. coli* BL21, refolded on a nickel NTA column, and purified by anion exchange (AIEX) chromatography (Supplementary Fig. 1A). All protein variants expressed and refolded in quantities similar to that of MMP-9_Cat_ WT and eluted at similar times.

Designs were tested for their ability to cleave MMP-9 substrate in an activity assay that measures the cleavage of a fluorogenic substrate peptide, resulting in fluorescence at 395 nm [31]. All four designs showed activity comparable to MMP-9_Cat_ WT, indicating that they were properly folded (data not shown). We next evaluated whether our variants were stable to self-cleavage by analyzing them with SDS-PAGE immediately after purification with AIEX. Gel electrophoresis analysis showed that Des1, Des3, and Des4 exhibited partial self-cleavage immediately after purification similar to that of MMP-9_Cat_ WT while Des2 did not exhibit self-cleavage (Supplementary Fig. 1B). We hence selected Des2 for further evaluation.

We next sought to compare the auto-cleavage activity of Des2 to that of MMP-9_Cat_ WT over a longer period of time. For this purpose, we incubated a 0.5µM sample of Des2 and MMP-9_Cat_ WT at 37 C for a maximum of seven days and analyzed the samples on SDS-PAGE as time progressed (Fig. 3A). Our data show that an auto-cleavage band appeared in the MMP-9_Cat_ WT sample already after 1 hour of incubation and the protein degraded almost completely after 7 days (Fig. 3A, top). In contrast, Des2 showed no auto-cleavage product even at the end of the 7-day period (Fig. 3A, bottom).

**Figure 3.**
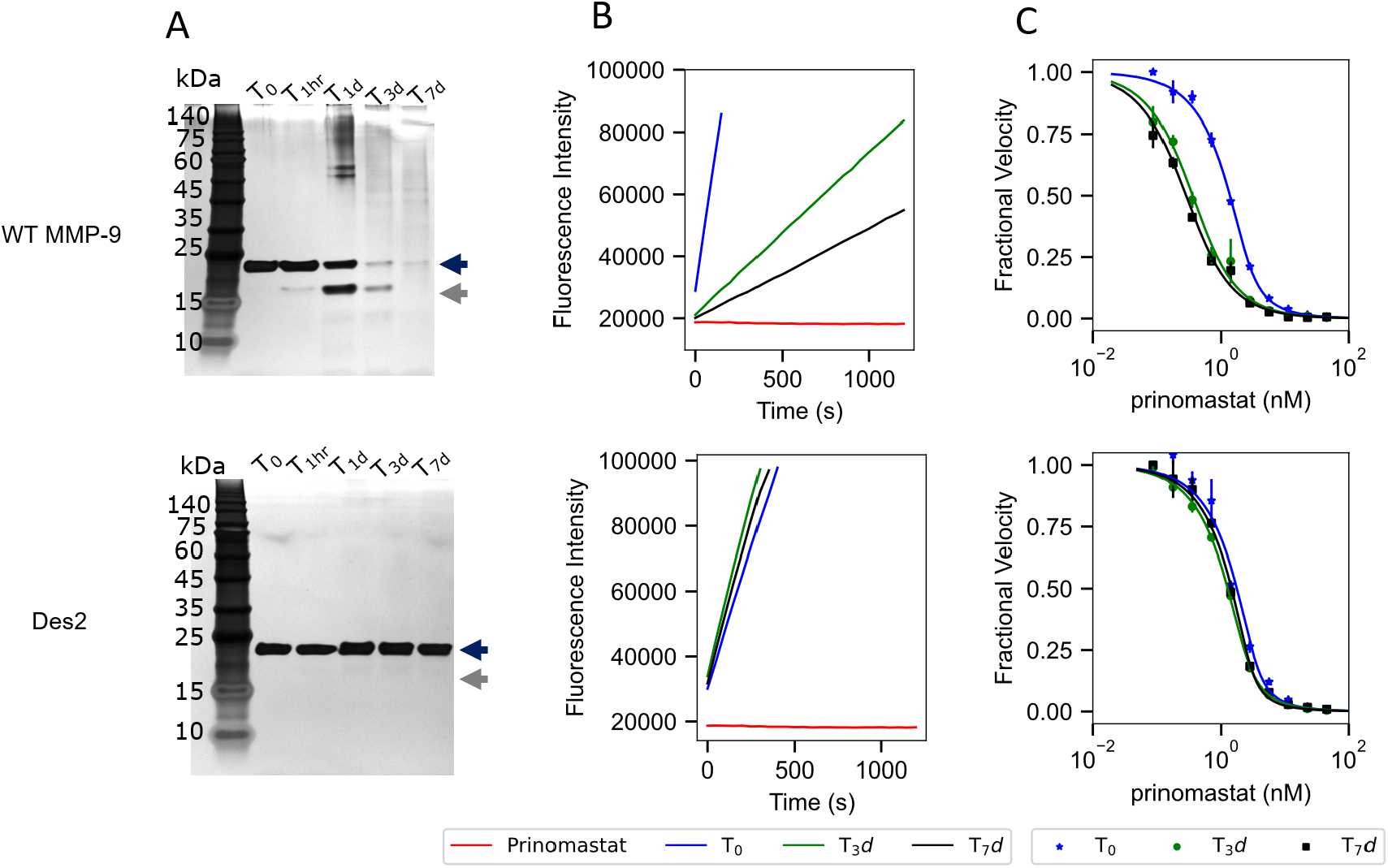
Evaluation of MMP-9_Cat_ WT and Des2 stability and enzyme activity at different time points. (**A**) Silver stained SDS-PAGE analysis showing full-length and cleavage products of MMP-9_Cat_ WT (top) and Des2 (bottom) after 37ºC at several time intervals, T = 0, T =1 hour (T_1hr_), T= 1 day (T_1d_), T= 3 days (T_3d_), and T= 7 days (T_7d_). (**B**) Enzyme activity of MMP-9_Cat_ WT (top) and Des2 (bottom) at T_0_, T_3d_, and T_7d_ using a fluorogenic peptide substrate. Substrate digestion was monitored by the appearance of a fluorescence signal at 395nm. (**C**) Inhibition of MMP-9_Cat_ WT (top) and Des2 (bottom) at T_0_, T_3d_, and T_7d_ using prinomastat as an inhibitor, using the same fluorogenic substrate as in B. K_i_^app^ was determined by fitting the data to Morrison’s equation (see equation 1 in Methods) [29].

We next measured the enzymatic activity of MMP-9_Cat_ WT and Des2 at three time points (T=0, T=3d, and T=7d) (Fig. 3B). In agreement with the SDS-PAGE results, MMP-9_Cat_ WT activity decreased with successive time points as more enzyme is cleaved, while Des2 maintained 100% activity at all time points. The same tendency was observed when evaluating enzyme’s inhibition by its strong active-site inhibitor, prinomastat (Fig. 3C, Table *2*). While the concentration of the active enzyme for MMP-9_Cat_ WT decreased as time progressed, it remained constant for Des2 (Fig. 3C, Table 2), attesting to the structural integrity of the enzyme’s active site in Des2 but not in MMP-9_Cat_WT after several days of incubation.

**Table 2.** Active Enzyme concentration and Ki^app^ for prinomastat of MMP-9_Cat_ WT and Des2 at different time points. The parameters are obtained by fitting Figure 3C to equation 1.

|  | T<sub>0</sub> |  | T<sub>1</sub> |  | T<sub>2</sub> |  |
| --- | --- | --- | --- | --- | --- | --- |
| | $K_i^{app}$, nM | [E], nM | $K_i^{app}$, nM | [E], nM | $K_i^{app}$, nM | [E], nM |
| MMP-9<sub>Cat</sub> WT | $0.30 \pm 0.04$ | $2.1 \pm 0.2$ | $0.22 \pm 0.02$ | $0.27 \pm 0.05$ | $0.16 \pm 0.03$ | $0.18 \pm 0.09$ |
| Des2 | $0.28 \pm 0.03$ | $2.9 \pm 0.4$ | $0.268 \pm 0.003$ | $2.00 \pm 0.03$ | $0.18 \pm 0.04$ | $2.5 \pm 0.3$ |

## Discussion

In this study, we present a novel method for designing protease enzymes that are active and stable to self-cleavage, which is based on the initial identification of the autocleavage site and subsequent computational design of non-autocleaving variants. The computational design protocol considered a combination of auto-degradation score, design energy, and the recapitulation of sequence and structural features found in other MMP homologs. Out of four tested variants, one designed variant, Des2, exhibited zero auto-cleavage and retained full catalytic activity up to 7 days of incubation at 37ºC, a remarkable stability for a proteolytic enzyme. Furthermore, Des2 bound an active-site MMP-9 inhibitor with the same Ki^app^ as MMP-9_Cat_ WT, indicating that the active site and the enzyme structural properties are unaltered.

In previous studies, inactive catalytic-site-mutants of MMP-9 were developed, by mutating the catalytic E402 to alanine or asparagine [25,26,32]. Corresponding residues were also mutated in other MMPs [33–37]. These mutants were considerably more stable than the WT MMP constructs and have been utilized to solve crystal structures of various MMPs [26,38]. However, such proteins remained inactive and could not be used in applications such as drug discovery. In contrast, we mutated only the identified self-cleavage site and introduced only two mutations, leaving the catalytic site unaltered. Our modeling with AlphaFold shows that Des2 exhibits an almost identical structure to MMP-9_Cat_ WT overall and especially in the region of the redesigned loop (Fig. 2D). Our engineered variant hence offers multiple advantages for studies that require working with MMP-9_Cat_ at high concentrations while preserving its catalytic activity. It is particularly suitable for discovery of active-site MMP-9 inhibitors, that are being conducted in various research laboratories and pharmaceutical companies.

Native enzymes including proteases are intrinsically unstable and frequently difficult to purify and work with. Computational design could be applied to significantly improve enzyme stability. Fleishman and co-workers devised a protocol that allows enzyme stabilization by introducing multiple mutations far from the active site observed in the enzyme evolutionary profile [39]. This strategy, however, achieves enzyme stabilization via multiple mutations and could not be applied to our project where we want to keep MMP-9_Cat_ WT sequence as close to WT as possible. Hence, we developed a different method that combines mass spectroscopy, self-cleavage and stability predictions to design out the self-cleavage site in MMP-9_Cat_ WT.

Although all four designed MMP-9_Cat_ WT variants were active, only Des2 exhibited stability to self-cleavage. This is consistent with its predicted self-cleavage score (Table 1), which is considerably better than that of MMP-9_Cat_ WT. Des1, Des3, and Des4 exhibit self-cleavage scores only slightly better than that of MMP-9_Cat_ WT. Thus, our results established the range of the Procleave score predictions that produce substantial changes experimentally and could be used to evaluate future self-cleavage-resistant variants.

In conclusion, we computationally designed an MMP-9_Cat_ variant that retained WT-like catalytic activity and binding to a small molecule inhibitor yet abolished undesired auto-degradation property. Our methodology could be applied to the redesign of other enzymes prone to self-cleavage, facilitating future studies of enzyme characterizations, structural determination, and drug design efforts.

## Methods

### Design of the non-degrading MMP-9_Cat_ variant

For energy calculations, single-position computational mutational scan and multi-position designs were built using Rosetta FastDesign [30], running 20 trajectories per position or multi-position design, and then taking the average score of the best 5. For these calculations, MMP-9_Cat_ crystal structure (PDB: 5TH6) with the pro-domain removed was relaxed using Rosetta FastRelax with SetupMetalsMover [40] and used as the input model in all runs. The backbone coordinates were constrained to the input model, and the option use_input_sidechains was used. Only neighbors within 6 Å to any of the 8 positions mutated in the mutational scan were allowed to be repacked. The structures of the designed proteins were visually compared to these of the MMP-9 homologs for sequence identity and rotamer conformation similarity.

For the predicted auto-cleavage score, we used the web server procleave developed for substrate-cutting prediction of MMP9_Cat_ [29], using as input the designed MMP9_Cat_ sequences.

### DNA cloning and Mutagenesis

The primers containing mutation (mutagenesis primers) for Des1 and Des2 were designed and purchased from Integrated DNA Technologies (IDT DNA, USA) and cloned into pET28 plasmids using the Transfer PCR protocol (TPCR) [41]. Gene blocks of MMP-9_Cat_ variants Des 3 and Des 4 were purchased from IDT DNA, USA. Primary PCR was carried out to generate the megaprimer, which was used to clone the WT MMP-9 pET28a plasmid using Restriction free Cloning [42]. All primers are summarized in Supplementary Table 1. PCR products were run on a 1% agarose gel to confirm the proper size of the expected PCR product (∼6000 bp). PCR products were DpnI digested to eliminate parental bacterial plasmid DNA. pET28a plasmids encoding MMP-9_Cat_ WT & variants were transformed into *E. coli* BL21 (DE3) cells and plated on LB agar plates containing the antibiotic kanamycin (KAN) incubated at 37ºC overnight. A single colony was inoculated in 10 mL LB containing kanamycin and grown at 30°C overnight, resulting in several colonies the following day. Five colonies were incubated at 37 °C overnight in 5mL LB media supplemented with 50µg/mL of kanamycin to ensure the desired mutagenesis. DNA was isolated using the Mini prep protocol by Geneaid kit (Geneaid, Taiwan) and measured using NanoDrop 1000 spectrophotometer. We confirmed the correct sequence of the mutants with sanger sequencing (HyLabs).

### Protein expression

MMP-9_Cat_ WT and four variants were expressed in expressed in BL21 (D3E3) cells in a pET 28A plasmid as previously described [43]. Cells were transferred to 1 liter of Liquid Broth (LB) supplemented with the same concentrations of kanamycin. This culture was grown at 30°C until an OD at 600 nm of 0.8-1.2 was reached. Protein expression was then induced with the addition of 1 mM isopropyl-D-1-thiogalactopyranoside (IPTG), and the culture was allowed to grow overnight at 30°C. The following day, cells were pelleted at 10000 rpm for 15 minutes in a SLA1500 or GSA rotor. The pellet was then frozen at -20°C until further processing. Next, 1-liter pellets were unified and suspended in 30 ml buffer B [25mM Tris, 0.5M NaCl, 0.5% v/v triton, pH 8.0) with lysozyme (100ug/ml). Finally, the solution was homogenized and incubated at 4°C for 2 hours. To wash the inclusion from contaminants, this solution was sonicated (VCX 750 Sonicator cat no. H-1006-2) at 60% amplitude with a large tip probe (13 mm) with 5-sec pulses and 5-sec rest for 2 min until the solution changed. Next, this solution was centrifuged at 12500 rpm for 20 minutes at 4°C with a SS34 rotor. Sonication and centrifugation were repeated three more times. Next, the inclusion was suspended in 50 ml solubilization buffer C (6M Urea, 25 mM Tris, pH 8.0) to solubilize the inclusion body pellets. The solution was then sonicated until it was homogenous.

### Protein Refolding

The protein renaturation process was done on a gravity column at 4ºC in the presence of 0.2% zwittergent. First, the MMP-9_Cat_ WT and variants (Des1, Des2, Des3, Des4) were incubated with the Ni^2+^-NTA agarose affinity column (PureCube cat no. 31103) for an hour. Then, the unbound protein was removed using a wash buffer (6M Urea, 50mM Tris, 30mM NaCl, and 30mM Imidazole; pH 7.5). Next, stepwise removal of Urea was done using Buffer G (4M Urea, 50mM Tris, 30mM NaCl, 5mM CaCl_2_, 20 µM ZnCl_2’_ pH7.5), Buffer H (2M Urea, 50mM Tris, 30mM NaCl, 5mM CaCl_2_, 2 µM ZnCl_2,_ pH 7.5) for 2hrs, and a refolding buffer (50mM Tris, 30mM NaCl, 5mM CaCl_2_, 20 µM ZnCl_2,_ pH 7.5) overnight. Finally, the proteins were eluted with Elution Buffer (50 mM Tris, 30 mM NaCl, 5mM CaCl_2_, and 250 mM imidazole, pH 7.5) and purified using Anion exchange chromatography (Hitrap Q HP, Cytiva). We further optimized this renaturation protocol by introducing another additive, a weak inhibitor of MMP-9, Acetohydroxamic acid (AHA) [35], at a concentration of 200 mM. The inhibitor was removed using Size exclusion (Hiload 16/60 Superdex 75). Protein concentration was determined by measuring the absorbance at 280 nm using a calculated extinction coefficient of 33920 M−1 cm−1.

### Enzymatic activity assay of MMP-9Cat and designs

The proteolytic activity of MMP-9_Cat_ WT, Des1, Des2, Des3, and Des4 was measured using a fluorogenic substrate Dnp-Pro-Leu-Gly-Leu-Trp-Ala-D-Arg-NH2 (Sigma-Aldridge, USA) in an enzyme activity buffer (50 mM HEPES, 0.1 M NaCl, 10 mM CaCl_2_, and 0.05% Brij 35, pH=7.5). Varying concentrations of prinomastat (AG3340, Sigma-Aldridge, USA) were incubated with MMP-9_Cat_ WT or variant at a final concentration of 2 nM under conditions similar to that described in [21]. Fluorescence resulting from cleavage of the fluorogenic substrate was measured at 395 nm, at least every 10 s for at least 5 min, immediately after adding fluorogenic substrate with irradiation at 325 nm on a Biotek Synergy H1 plate reader (BioTek, Winooski, VT, USA). Normalized velocities were fitted in MATLAB Curve Fitting Toolbox (MathWorks) to the Morisson equation:

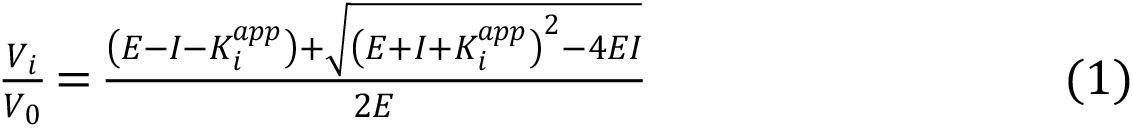

where Vi is enzyme velocity at inhibitor concentration I, V_0_ is the enzyme velocity in the absence of an inhibitor, E is the active enzyme concentration, and K_iapp_ is an apparent inhibitory constant. At least three assays were performed and average values were calculated.

### Self-cleavage Assay

Samples of pure MMP9_Cat_ WT and variants at concentration of 0.5µM were incubated in assay buffer (50mM Tris, 100mM NaCl, 5mM CaCl2, 0.05% Brij35) at 37 °C for several time points (T0, T1hr, T1d, T3d, T7d) and frozen at -80. Afterward, the auto-cleavage profile of the protein was analyzed on a silver-stained SDS-PAGE gel.

### LC-MS/MS data analysis

SDS-PAGE gels were thoroughly stained using Coomassie blue and de-stained using the de-staining solution (H_2_O, methanol, and acetic acid in a ratio of 50/40/10 (v/v/v)). MMP-9_Cat_ WT full-length and degradation product bands were excised and chopped into small cubes. The gel pieces were destained (25 mM Tris-HCl, pH 8.0 containing and acetonitrile (ACN)), treated with 10 mM Dithiothreitol (DTT) (Sigma Chem. Co.) and 55 mM iodoacetamide (Sigma Chem. Co.) and then digested with trypsin (Mass Spectrometry grade, from Promega Corp., Madison, WI, USA) overnight. The released peptides were extracted, desalted, and eluted using 80% ACN and 0.1% formic acid. 0.25 µg of eluted peptides was used for each sample to obtain the raw data. Mass spectrometry raw data was processed and searched using the Trans-Proteomic Pipeline (TPP) 6.0.0 “OmegaBlock” [44]. Searches were performed using Comet (2020.01 rev. 1) and high-resolution settings. They include ‘semi-tryptic’ cleavage specificity and oxidation of methionine and protein N-terminal acetylation as variable modifications. All searches were conducted against e. coli protein sequences downloaded from UniProt supplemented with the sequence of MMP-9_Cat_. Following the database search the putative self-cleavage sites generated by MMP-9 were mapped to the sequence using <u>PeptideMatcher</u> script and the P1 positions of cleavage sites were assigned with a value calculated from integrated MS/MS count of all the corresponding peptides.

## Data Availability

Mass Spectroscopy data is uploaded as Supplementary data.

## Author contributions

Conceptualization: A.B. Contributed equally: A.B. and S.O. Methodology: A.B., J.S.M., and S.O. Software: A.B. Formal analysis: A.B., J.S.M., O.K., S.O., and T.L. Investigation: S.O. Writing: A.B., J.S.M., and S.O. with contributions from all authors. Supervision J.S.M. and O.K. Funding acquisition: J.S.M. and O.K.

