## Supplementary Figure 1 for "Computational design of Matrix Metalloprotenaise-9 (MMP-9) resistant to auto-cleavage"

**Supplementary Figure 1: Purification and stability analysis of MMP-9$_{Cat}$ WT and variants.** A) Anion exchange chromatograms of Des1, Des2, Des3, Des4, and MMP-9$_{Cat}$ WT. All variants elute at ~300 mM NaCl. B) SDS-PAGE gel analysis of Des1, Des2, Des3, Des4, and MMP-9$_{Cat}$WT immediately after refolding (refolded) and after AIEX purification. The band around 19.6 kDa corresponds to the full-length MMP-9$_{Cat}$WT, while the band around 16 kDa corresponds to the product after self-cleavage.

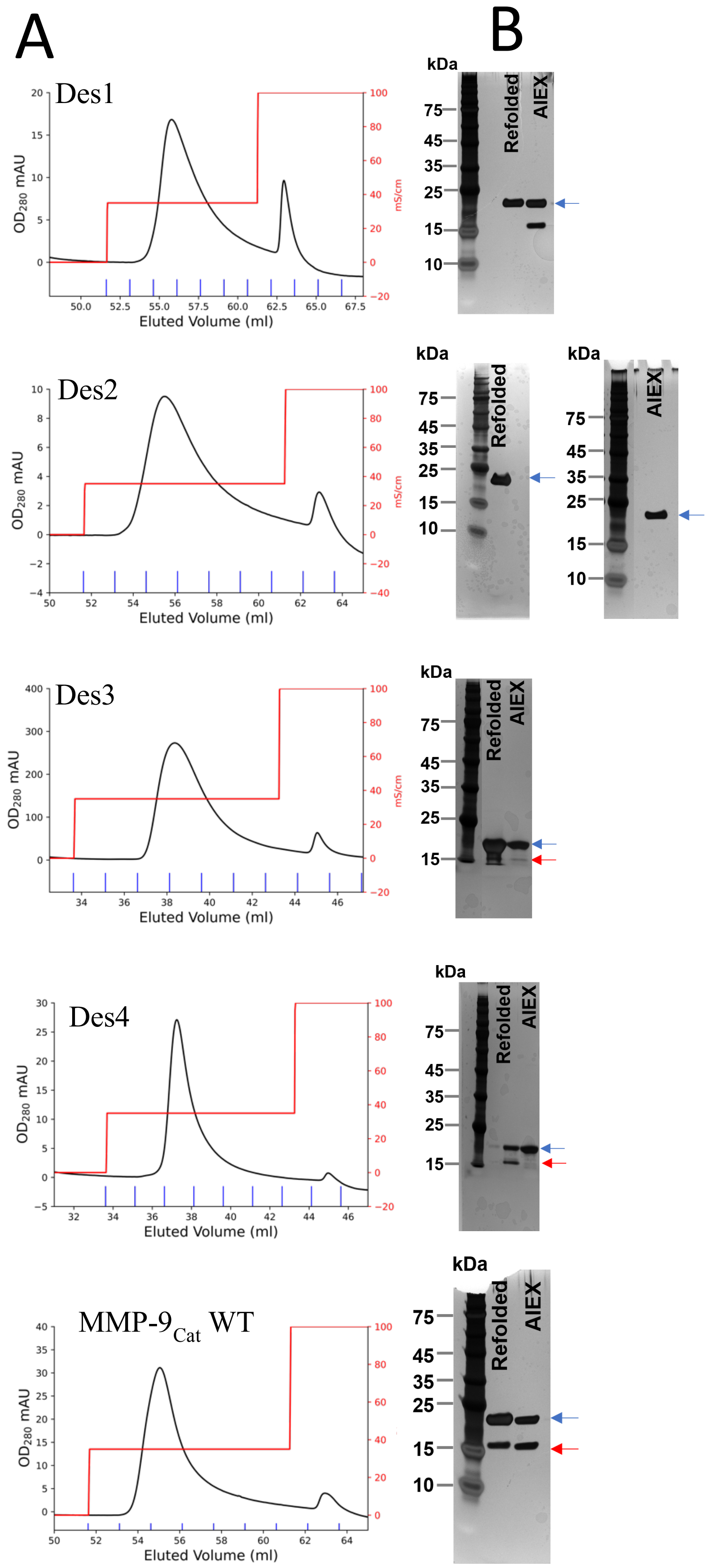

**Supplementary table 1**. Gene construct for MMP-9$_{Cat}$ variants.
General forward: GGCTTCCAGACCTTTGAAGGCGACTTGAAGTGGCATC

General Reverse: GCCGTTCACATCATCCTTATGCAAAGGCGGCCC

| Variant/Mutation | Reverse primer/gene block |
|---|---|
| DES1 (DES414D) | GTACATCAAGGCTTCGGGatcTGACGAATGATCCAGTCCC |
| DES2(DES414D & P415T) | CATCGGGTACATCAAGGCTTCggtgttTGACGAATGATCCAGTCCCA GTG |
| DES3 (L147I, A400V, L418I) | GGCTTCCAGACCTTTGAAGGCGACTTGAAGTGGCATCACCATAATA TTACCTACTGGATTCAAAA<br>CTATTCGGAGGACCTGCCGCGTGCCGTGATTGACGACGCGTTCGC GCGCGCGTTTGCCgtgTGGT<br>CAGCGGTTACTCCGCTGACTTTCACTCGCGTTTATAGCCGCGACGC CGATATCGTAATTCAGTTT<br>GGAGTTGCAGAACATGGTGATGGCTATCCTTTTGACGGTAAAGAT GGTCTGTTAGCACACGCGTT<br>TCCGCCTGGGCCGGGTATTCAGGGTGATGCACATTTTGATGATGAT GAACTGTGGTCTCTGGGCA<br>AAGGCCAGGGATATTCACTGTTTTTGGTTGCCgtgCATGAATTTGG GCATGCACTGGGACTGGAT<br>CATTCGTCAGTCCCCGAAGCCattATGTACCCGATGTATCGTTTCAC GGAAGGGCCGCCTTTGCA<br>TAAGGATGATGTGAACGGC |
| DES4(L147I, A400V, A417S, L418I) | GGCTTCCAGACCTTTGAAGGCGACTTGAAGTGGCATCACCATAATA TTACCTACTGGATTCAAAA<br>CTATTCGGAGGACCTGCCGCGTGCCGTGATTGACGACGCGTTCGC GCGCGCGTTTGCCgtgTGGT<br>CAGCGGTTACTCCGCTGACTTTCACTCGCGTTTATAGCCGCGACGC CGATATCGTAATTCAGTTT<br>GGAGTTGCAGAACATGGTGATGGCTATCCTTTTGACGGTAAAGAT GGTCTGTTAGCACACGCGTT |

| | |
|---|---|
| | TCCGCCTGGGCCGGGTATTCAGGGTGATGCACATTTTGATGATGATGAACTGTGGTCTCTGGGCA<br><br>AAGGCCAGGGATATTCACTGTTTTTGGTTGCCgtgCATGAATTTGGGCATGCACTGGGACTGGAT<br><br>CATTCGTCAGTCCCCGAAagcattATGTACCCGATGTATCGTTTCACGGAAGGGCCGCCTTTGCA<br><br>TAAGGATGATGTGAACGGC |
